# Interplay between *Pitx2* and *Pax7* temporally governs specification of extraocular muscle progenitors

**DOI:** 10.1101/2023.08.24.554745

**Authors:** Mao Kuriki, Glenda Comai, Shahragim Tajbakhsh

## Abstract

Gene regulatory networks that act upstream of skeletal muscle fate determinants are distinct in different anatomical locations. Despite recent efforts, a clear understanding of the cascade of events underlying the emergence and maintenance of the stem cell pool in specific muscle groups remains unresolved and debated. Here, we invalidated *Pitx2* with multiple *Cre*-driver mice prenatally, postnatally, and during lineage progression and showed that this gene becomes progressively dispensable for specification and maintenance of the extraocular muscle (EOM) stem cell pool, yet it is the major EOM upstream regulator during early development. Moreover, constitutive inactivation of *Pax7* postnatally showed a greater loss of muscle stem cells in the EOM compared to the limb, pointing to a relay between *Pitx2*, *Myf5* and *Pax7* for maintenance of the EOM stem cells. Further, we demonstrate that EOM stem cells adopt a quiescent state earlier that those in limb muscles and do not spontaneously re-enter in proliferation in the adult as previously suggested, yet EOMs have a significantly higher content of Pax7+ muscle stem cells per area pre- and post-natally. This unique feature could result from different dynamics of lineage progression *in vivo*, given the lower fraction of committed and differentiating EOM myoblasts. Finally, significantly less MuSCs are present in EOM compared to the limb in the *mdx* mouse model for Duchenne muscular dystrophy. Overall, our study provides a comprehensive in vivo characterization of muscle stem cell heterogeneity along the body axis and brings further insights into the unusual sparing of EOM during muscular dystrophy.

## INTRODUCTION

Skeletal muscle satellite (stem) cells (MuSCs) arise during late foetal development, and they contribute to postnatal growth of muscle fibers. They are the main players for reconstituting muscle fibres during regeneration. Interestingly, this population of stem cells emerges in an asynchronous manner in different anatomical locations and it exhibits an unusual diversity in developmental origins and genetic regulation depending on the muscle group in which it resides (Comai et al., 2014; Grimaldi et al., 2022; Grimaldi and Tajbakhsh, 2021). Understanding the physiological mechanisms controlling the development of different muscle groups, and how they are selectively deregulated or spared during disease and ageing, is crucial for devising therapeutic strategies particularly for a wide range of myopathies (Emery, 2002; Porter and Greaves, 2003). Extraocular muscles (EOMs) are known to be spared during ageing and in Duchenne, Becker, and some limb girdle muscular dystrophies (Emery, 2002; Kallestad 2011; Porter et al., 2003). While the underlying causes for their sparing are unclear, intrinsic differences in EOM MuSCs have been proposed as contributing factors (Di Girolamo et al., 2023; Evano et al., 2020; Randolph and Pavlath, 2015). Yet, the precise mechanisms governing the emergence and maintenance of these and other cranial MuSCs remain obscure.

Skeletal muscle cell commitment and differentiation throughout the body is regulated by basic helix-loop-helix myogenic regulatory factors (MRFs) comprising Myf5, Mrf4, Myod and Myogenin (Myog). Mouse knockout models have established a genetic hierarchy where *Myf5*, *Mrf4* and *Myod* control commitment and proliferation of myogenic progenitors, and *Myod, Mrf4* and *Myog* are involved in terminal differentiation (Comai and Tajbakhsh, 2014; Kassar-Duchossoy et al., 2004; Rudnicki et al., 1993). Despite this uniformity in the acquisition of myogenic cell fate, an underlying heterogeneity characterises the embryonic cells that emerge in different anatomical locations. In the trunk, the paralogous transcription factors *Pax3* (Paired Box 3) and *Pax7* (Paired Box 7) regulate the emergence of muscle stem (Pax3+ or Pax7+) and committed (Myf5+, Mrf4+ or Myod+) cells (Buckingham and Relaix, 2015; Comai and Tajbakhsh, 2014; Relaix and Zammit, 2012). Notably, somite-derived myogenic cells are eventually lost by apoptosis from mid-embryogenesis after myogenesis has initiated in *Pax3;Pax7* double-mutant mice (Relaix et al., 2005). In contrast, craniofacial muscles such as EOMs, facial, mastication and subsets of neck muscles, are derived from cranial mesoderm and do not express *Pax3* (Grimaldi and Tajbakhsh, 2021; Schubert et al., 2019; Tzahor, 2015; Ziermann et al., 2018). The early head mesoderm harbors instead a complementary set of upstream markers including *Tbx1*, *Pitx2* and the *Six* gene family (Harel et al., 2009; Kelly et al., 2004; Sambasivan et al., 2009; Wurmser et al., 2023). Notably, *Tbx1* (T-Box Transcription Factor 1), is an upstream regulator for the facial, jaw, neck, laryngeal and esophagus muscles, but not EOM (Kelly et al., 2004; Lescroart et al., 2022; Sambasivan et al., 2011; Schubert et al., 2019). Instead, *Pitx2* (Paired Like Homeodomain 2), which is also expressed in head muscles, plays a critical role as an upstream regulator of EOM development, as these muscles are lacking in *Pitx2* null mice (Gage et al., 1999; Kitamura et al., 1999), with phenotypes being dependent on the gene dosage (Diehl et al., 2006) and timing of deletion (Zacharias et al., 2011).

In contrast to the diversity in upstream regulators, *Pax7* marks all adult MuSCs and their ancestors from mid-embryogenesis (Comai and Tajbakhsh, 2014; Kassar-Duchossoy et al., 2005; Relaix et al., 2005; Seale et al., 2000). In the trunk, Pax7+ myogenic cells are primed by prior *Pax3* expression during embryonic development (Hutcheson et al., 2009). However, the mechanisms that govern the emergence and maintenance of cranial myogenic cells are poorly understood. As *Pax3* is absent in the EOMs, and *Pax7* is expressed after the MRFs (Horst et al., 2006; Nogueira et al., 2015), it is possible that *Pitx2* or other transcription factors such as the *Six* family members (Maire et al., 2020; Wurmser et al., 2023) specify the Pax7+ population in this location. Yet, as no lineage-specific deletion of *Pitx2* has been performed in EOM MuSCs, it is unclear: i) to what extent *Pitx2* is required temporally for emergence and maintenance of the MuSC population, and its relative function compared to the MRFs and Pax7; ii) if *Pitx2* is continuously required for maintenance of the adult stem cell population. Further, global deletion of *Pax7* in trunk and limb muscles leads to an extensive loss of MuSCs during early postnatal development (Kuang et al., 2006; Lepper et al., 2009; Oustanina et al., 2004; Relaix et al., 2006; Seale et al., 2000). Deletion of *Pax7* in adult MuSCs results in reduced MuSC numbers and impaired regeneration following muscle injury (Gunther et al., 2013; von Maltzahn et al., 2013). While *Pax7* is dispensable for EOM formation during embryonic development (Zacharias et al., 2011), the presence of MuSCs in the EOM of postnatal *Pax7* null mice and the relative roles of *Pax7* and *Pitx2* in maintaining EOM cells remain unresolved. Finally, several reports showed that *Pitx2* is expressed in postnatal EOM MuSCs at higher expression levels than in limb muscles (Di Girolamo et al., 2023; Evano et al., 2020; Taglietti et al., 2023). As high levels of *Pitx2* are retained in dystrophic and ageing mouse EOM (Hebert et al., 2013; Taglietti et al., 2023) and overexpression of *Pitx2* was reported to enhance the regenerative potential of dystrophic skeletal MuSCs (Vallejo et al., 2018), *Pitx2* could potentially contribute to the sparing of EOMs in muscular dystrophies. However, the functional role of *Pitx2* in EOM MuSCs in *mdx* mice has not been formally evaluated.

In this study, we examined the spatiotemporal functional roles of *Pitx2* and *Pax7* in the emergence and maintenance of the EOM progenitor population using combinations of genetically modified and lineage traced mice. Our study provides a qualitative and quantitative analysis of myogenesis from embryonic to postnatal stages in vivo and identifies differential cycling states, dynamics of commitment, and establishment of stem cell quiescence in EOM compared to limb muscles. We show that *Pitx2* is critically required for establishing EOM prior to *Pax7* expression but is subsequently dispensable. Moreover, our findings clarify the behavior of EOM MuSC behavior in *mdx* mice thus providing valuable developmental and clinical insights.

## RESULTS

### Differential temporal dynamics of EOM and limb myogenic cells during development and postnatally

Previous studies suggested that EOM have more MuSCs per area and per tissue weight than limb muscles such as the *Tibialis anterior* (TA) (Keefe et al., 2015; Verma et al., 2017). To determine if the higher density of EOM MuSCs *in vivo* results from a longer proliferative phase of EOM Pax7+ cells during embryonic and postnatal development, we performed co-immunostaining of Pax7 and the proliferation marker Ki67 on tissues sections from EOM and TA from E (embryonic day) 14.5 to 4 months postnatally (Fig 1A, 1B). As expected, the total numbers of Ki67+/Pax7+ cells were progressively reduced during postnatal growth in both muscles. Yet, EOM MuSCs stopped proliferating by P (postnatal day) 14, whereas proliferating Pax7+ MuSCs continued to be observed in TA muscle at P20 (Fig. 1C). Thus, EOM Pax7+ cells exited the cell cycle earlier than those in the TA. In the TA, cycling Pax7+/Ki67+ cells persisted in pre-pubertal muscle (P28) before quiescence was achieved between 7-8 weeks (Gattazzo et al., 2020) (Fig. 1C). Notably, the density of EOM Pax7+ MuSCs remained higher in the orbital layer, probably due to the smaller fibre size and less myoblast fusion events during the growth phase compared to the global layer (Fig. S1A). Altogether, these data show that the relatively higher number of Pax7+ cells per area in the EOM cannot be accounted for by a longer proliferative phase. Of note, as *Pitx2* was expressed in the majority of EOM Pax7+ cells throughout all stages examined (80-90%; Fig. 1D, 1E), its expression alone does not seem account for the higher proliferative status of this population.

**Figure 1.**
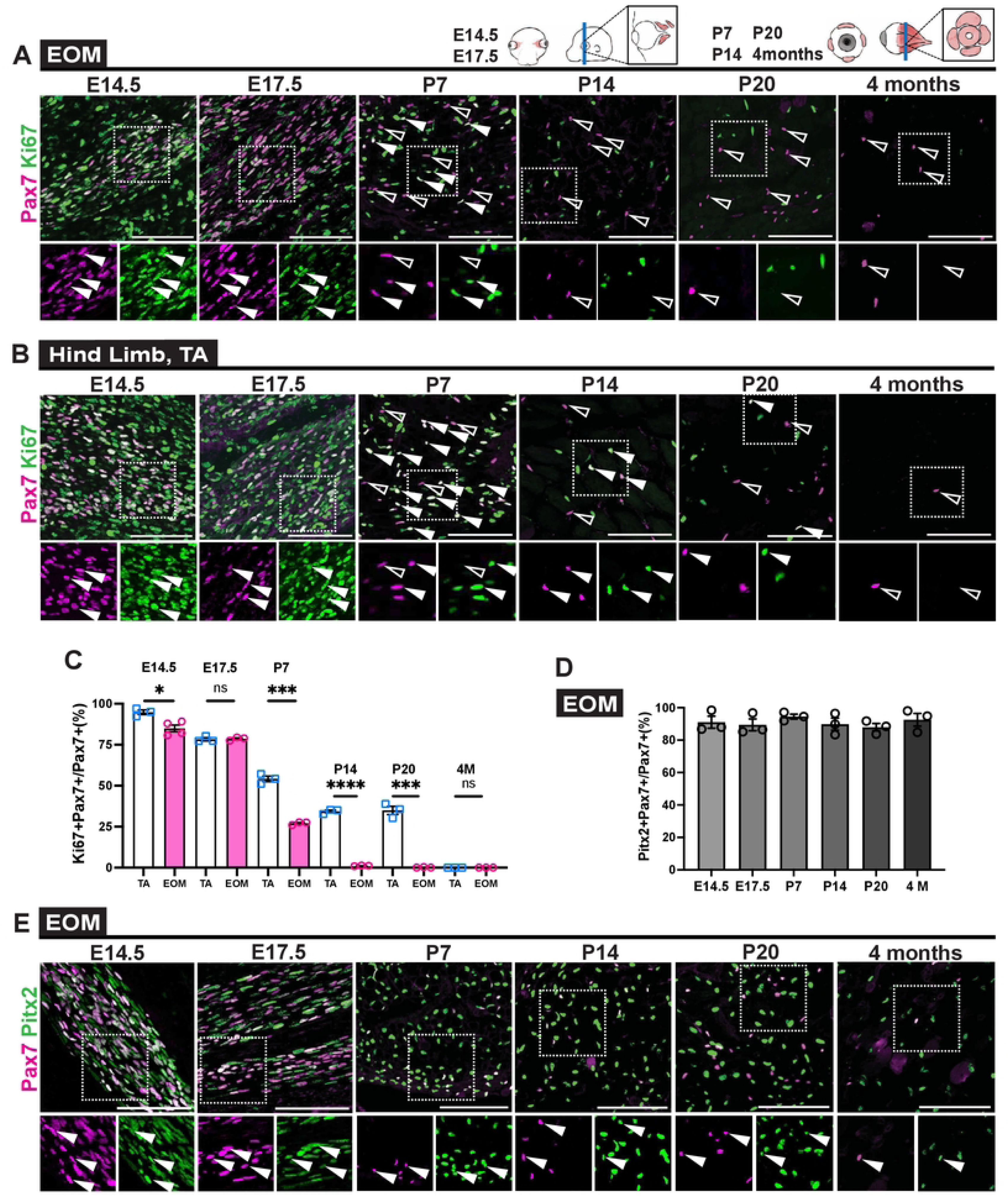
Postnatal EOM myogenic cells exit the cell cycle earlier than those in TA muscle. **(A-B)** Immunostaining of EOM and TA sections for Pax7 (magenta) and Ki67 (green) at indicated stages from E14.5 to 4 months. White arrowheads point to Ki67+/Pax7+ cells, open arrowheads to Ki67-/Pax7+ cells. Bottom panels, higher magnification views of the area delimited with dots. EOM are composed of 4 rectus (medial, lateral, dorsal, ventral) and 2 oblique (superior, inferior) muscles (Comai et al., 2020; Suzuki et al., 2016). **(C)** Percentage of Ki67+/Pax7+ cells in EOM and TA muscles at each stage (n=3 per stage). **(D)** Percentage of Pitx2+/Pax7+ cells over total Pax7+ stem cells at indicated stages from E14.5 to 4 months (4M). Note that *Pitx2* is expressed expresses in about 80∼90% of myogenic cells in EOM throughout all stages (n=3 each stage). **(E)** Immunostaining for Pax7(magenta) and Pitx2(green) on EOM sections at indicated stages. White arrowheads point to Pitx2+/Pax7+ cells. Bottom panels, higher magnification views of the area delimited with dots. Scale bars: 100μm (A), (B) and (E). Error bars represent SEM. ns, non-significant, *P<0.05, ***P<0.001, ****P<0.0001.

### *Pax7* is critical for maintenance of EOM MuSC pool postnatally

To investigate the role of *Pax7* in EOM MuSCs compared to limb muscles, we examined *Pax7^nGFP/nGFP^* knock-out mice (*Pax7* KO) (Sambasivan et al., 2013; Sambasivan et al., 2009) where *Pax7*-null and heterozygous cells are GFP+ (Fig. 2A, 2B). At P20, the number of GFP+ cells/area was significantly reduced both in EOM and TA of *Pax7* KO mice compared to heterozygous controls, although to a greater extent in the EOM (Fig. 2C). Assessment of the number of GFP+ cells/100 fibres revealed similar reductions in reduction in the EOM and TA of *Pax7* KO mice compared to heterozygous controls (Fig. S2 A-C). Interestingly, *Pax7* KO mice showed a reduction in GFP+ cells principally in the orbital (outer) EOM layer, yet this was relatively unchanged in the global layer, thereby pointing to location specific requirements for *Pax7* in this muscle (Fig. S2 A, B).

**Figure 2.**
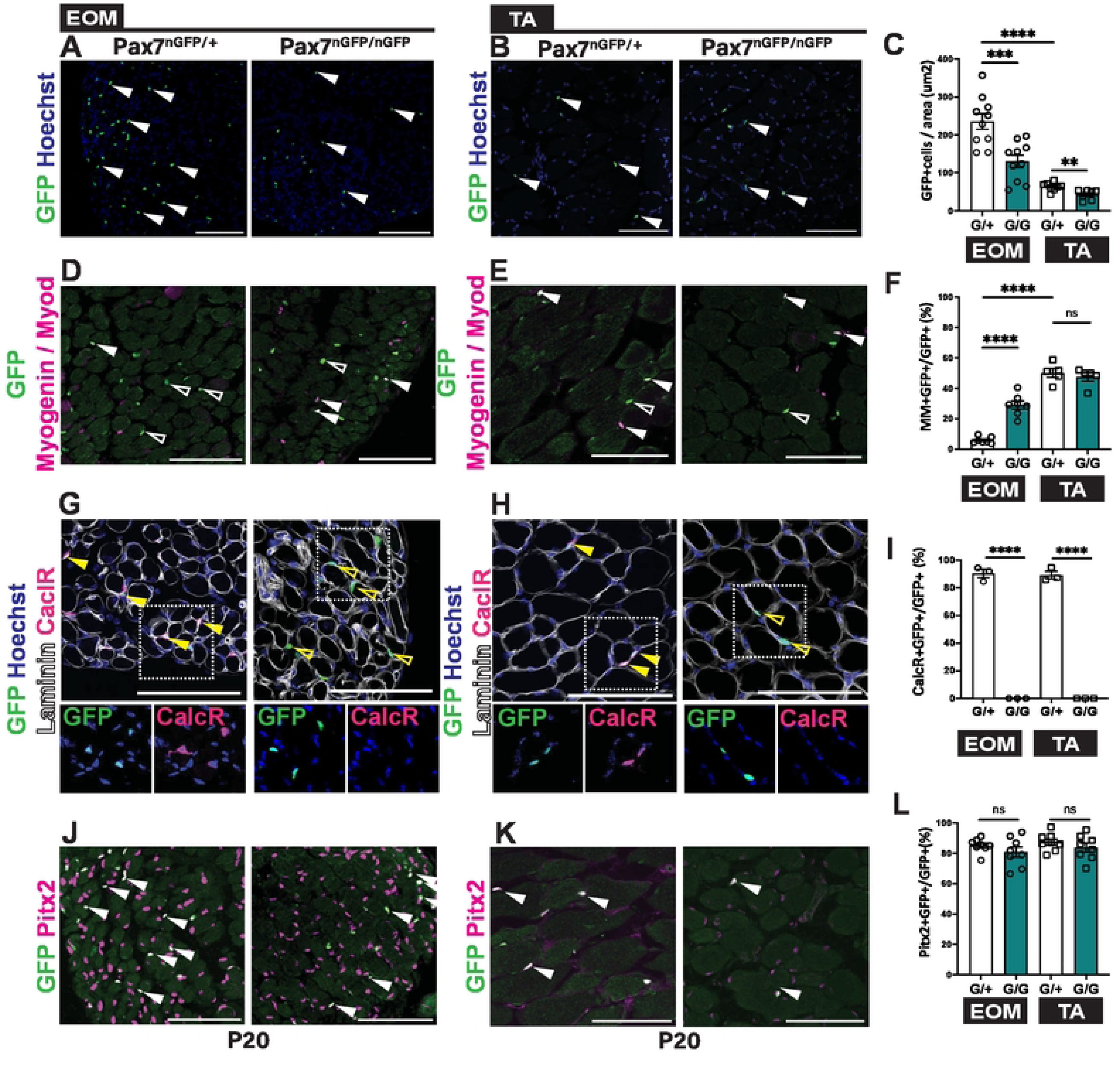
*Pax7* is critical for the maintenance of EOM MuSC pool postnatally. **(A,B)** Immunostaining of EOM and TA sections from *Pax7^nGFP/+^* and *Pax7^nGFP/nGFP^* P20 mice for GFP (green). White arrowheads indicate GFP+ cells. **(C)** Number of GFP+ cells per area in EOM and TA sections from the immunostaining in (A-B). (EOM n=10, TA n=8). **(D,E)** Immunostaining of EOM and TA sections from *Pax7^nGFP/+^* and *Pax7^nGFP/nGFP^* P20 mice for GFP (green), together with Myod/Myogenin (MM) (magenta). White arrowheads indicate MM+/GFP+ cells; open arrowheads MM-/GFP+ cells. **(F)** Percentage of MM+/GFP+ cells over total GFP+ cells in EOM and TA sections from immunostaining in (D,E) (EOM n=7, TA n=5). **(G,H)** Immunostaining of EOM and TA sections from *Pax7^nGFP/+^* and *Pax7^nGFP/nGFP^* P20 mice for GFP (green), Calcitonin receptor (CalcR) (magenta) and Laminin (white). Yellow arrowheads indicate CalcR+/GFP+ cells; yellow opened arrowheads indicate CalcR-/GFP+ cells. Higher magnification views of the area delimited with dots. **(I)** Percentage of CalcR+/GFP+ cells in GFP+ cells in EOM and TA sections from immunostaining in (G,H) (n=3 each). **(J,K)** Immunostaining of EOM and TA sections from *Pax7^nGFP/+^* and *Pax7^nGFP/nGFP^* P20 mice for GFP (green), together with Pitx2 (magenta). White arrow heads indicate Pitx2+/GFP+ cells. **(L)** Percentage of Pitx2+/GFP+ cells over total GFP+ cells in EOM and TA sections from immunostaining in (J,K) (EOM n=8, TA n=8). G/G: *Pax7^nGFP/nGFP^* G/+: *Pax7^nGFP/+^*. Scale bars: 200μm (A) and (B), 100μm (D,E,G,H,J,K). Error bars represent SEM. ns, non-significant, **P<0.01, ***P<0.001, ****P<0.0001.

Next, we addressed the mechanisms underlying MuSC loss in *Pax7* null EOM. Progressive loss of limb and trunk MuSCs in newborn constitutive *Pax7* mutants was suggested to be due to a proliferation defect and cell death (Oustanina et al., 2004; Relaix et al., 2005). Moreover, a greater propensity to differentiate and fuse into myofibres was reported for postnatal *Pax7* mutant MuSCs *in vivo* (Lepper et al., 2009). Similarly, adult limb and trunk *Pax7* mutant cells displayed reduced proliferation and increased differentiation *in vitro* (Gunther et al., 2013; von Maltzahn et al., 2013). We performed immunostaining for Myod (commitment marker) and Myogenin (Myog, differentiation marker) in *Pax7* EOM and TA muscles of *Pax7* KO and control mice (Fig. 2D, E). At P20, a significantly higher fraction of Myod+/Myog+ (MM+) cells in the GFP+ population was observed in *Pax7* KO EOM (30% in KO vs 5.8% in control; Fig. 2F), indicative of depletion of the stem cell population. This difference was not notable for TA muscle at P20. We then examined EOM and TA muscle of *Pax7* KO E14.5 embryos to assess the role of *Pax7* in maintenance of myogenic progenitors in the foetus (Fig. S2D-L). At E14.5, the number of Pax7+ cells per area was similar between EOM and TA of *Pax7* KO embryos (Fig. S2F). However, the fraction of MM+/GFP+ cells was already higher in the EOM of *Pax7* KO embryos compared to controls (Fig. S2I). Notably though, the control TA had ∼9-fold and ∼2-fold more Myod+/Myog+ cells in the GFP+ population compared to the EOM at P20 (Fig. 2F) and E14.5 (Fig. S2I), respectively, indicating a more rapid transition from stem to committed and differentiated cells in vivo in normal conditions. Evaluation of the expression of Calcitonin receptor (CalcR), which is a marker of postnatal quiescent MuSCs (Fukada et al., 2007), showed that >90% of GFP+ cells in the *Pax7* heterozygous were positive for this marker at P20 as expected, whereas all GFP+ cells in the *Pax7* KO were CalcR-negative in both EOM and TA muscles, suggesting that in both muscles this residual MuSC population is perturbed (Fig. 2G-I). Altogether, we conclude that Pax7 is required for maintenance of the stem cells population of the EOM, and that invalidation of this gene results in a faster loss of the upstream population compared to the limb.

Given that *Pitx2* is a major upstream regulator of the EOM lineage (Sambasivan et al., 2009; Zacharias et al., 2011), we assessed whether Pitx2+ cells would expand in the *Pax7* KO. Co-immunostaining for Pax7 and Pitx2 in *Pax7* KO mice at embryonic and adult stages showed that the number of Pitx2+/GFP+ cells were similar in EOM as well as TA in the *Pax7* KO and control mice (Fig. 2J-L and S2J-L). Moreover, unlike other skeletal muscles, EOMs were reported to contain another population of interstitial stem cells that are Pax7-/Pitx2+ (Hebert et al., 2013). Thus, we examined this interstitial population in mutant versus control EOMs and showed similar proportions of Pitx2+/GFP-cells between two (Fig. S2M). Overall, these results suggest a lack of compensation by Pitx2 in the EOM lineage of *Pax7* KO mice.

### Deletion of *Pitx2* in Myf5+ cells impair establishment of EOM and emergence of Pax7+ cells

Throughout the body, the long-term stem cell population (Pax3+/Pax7+ in the trunk; Pax7+ in head) emerges only after an anlage of differentiated cells has been established during mid-embryogenesis (Comai and Tajbakhsh, 2014). Cranial muscles are specified temporally after those in the trunk (Bothe et al., 2011; Gage et al., 1999; Horst et al., 2006; Nogueira et al., 2015; Sambasivan et al., 2009) and Pax7+ stem cells appear at later stages during development, after expression of the myogenic markers Myf5 and to a lesser extent Myod (Nogueira et al., 2015). However, it is unclear if expression of the MRFs is a prerequisite for the emergence of the Pax7+ cells themselves.

Therefore, we performed whole mount immunofluorescence (WMIF) analysis on *Myf5^nlacZ/+^* mice, where β-galactosidase serves as a proxy for *Myf5* expression (Tajbakhsh et al., 1996) (Fig. 3A-C). In the EOM primordium, *Pitx2* was expressed broadly, including in periocular mesenchyme and connective tissue cells (Comai et al., 2020; Gage et al., 1999). WMIF showed that virtually all Myf5+ cells expressed *Pitx2* in the EOM anlage throughout the stages analysed (Fig. 3B). In addition, we observed Pax7+ expression from E11.75, only in Myf5+ cells (Fig. 3C). EOM Pax7+ cells then rapidly increased in numbers and together with Pax7 protein levels from E11.75 (Fig. 3B, 3C).

**Figure 3.**
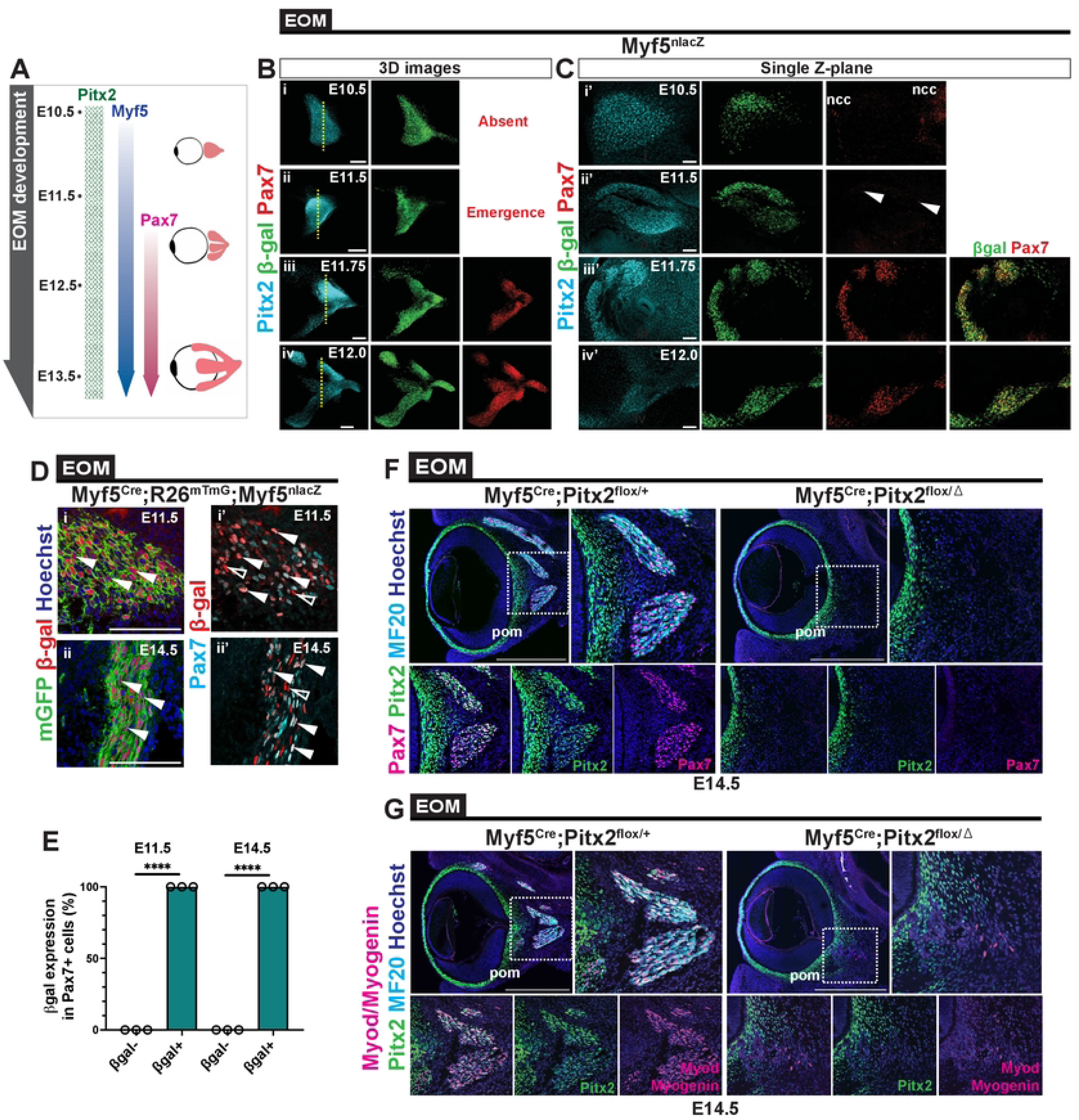
Deletion of *Pitx2* in the *Myf5* positive cells impairs EOM establishment. **(A)** Scheme summarizing the timing of expression of *Pitx2*, *Myf5* and *Pax7* during EOM development. **(B)** Whole-mount immunostaining of *Myf5^nlacZ^* at (i) E10.5, (ii) E11.5, (iii) E11.75, (iv) E12.0 for Pax7, β-gal and Pitx2. EOMs were segmented from adjacent head structures and 3D-reconstructed in Imaris (Bitplane). Yellow dashed lines indicate Z-section level shown in (C). **(C)** Single Z-section of the respective segmented volumes at (i’) E10.5, (ii’) E11.5, (iii’) E11.75, (iv’) E12.0 from (B). ncc, neural crest cells. Only few cells express Pax7 at E11.5 (white arrowheads). **(D)** Immunostaining of EOM sections from *Myf5^Cre^;R26^mTmG^: Myf5^nlacZ^* embryos at E11.5 (i, i’) and E14.5 (ii, ii’) for Pax7 (cyan), β-gal (red) and GFP (green). White arrowheads in (i) and (ii) indicate mGFP+/β-gal+ cells. White arrowheads in (i’) and (ii’) indicate Pax7+/β-gal+ cells. White open arrowheads point to Pax7-/β-gal+ cells. **(E)** Percentage of β-gal+ and negative cell over total Pax7+ cells in EOM at E11.5 and E14.5 from the immunostaining in (D) (n=3 each). **(F)** Immunostaining of E14.5 EOM sections from *Myf5^Cre^;Pitx2^flox/+^* (control) and *Myf5^Cre^;Pitx2^flox/^* ^Δ^ (KO) at for Pax7 (magenta), Pitx2 (green) and MF20 (cyan). Bottom panels, higher magnification views of the area delimited with dots. Periocular mesenchyme (pom). **(G)** Immunostaining of E14.5 EOM sections from *Myf5^Cre^;Pitx2^flox/+^* (control) and *Myf5^Cre^;Pitx2^flox/^* ^Δ^ (KO) for Myod/Myogenin (MM) (magenta), Pitx2 (green) and MF20 (cyan). Bottom panels, higher magnification views of the area delimited with dots. Periocular mesenchyme (pom). Scale bars: 100μm (D) and (B), 40μm (C)(i’) and (C)(ii’), 50μm (C)(iii’) and (C)(iv’) 500μm (F) and (G). Error bars represent SEM. ****P<0.0001.

To examine further the emergence of EOM MuSCs, we performed lineage tracing of Pax7+ cells originating from Myf5+ cells (Haldar et al., 2007) using the *Myf5^Cre^;R26^mTmG^;Myf5^nlacZ^* genetic combination (Fig. 3D). This genetic strategy allowed us to first assess the number of Pax7+ cells with a history of *Myf5*-expression (membrane GFP+), then to detect with high sensitivity, contemporary *Myf5*-expressing cells (β-gal+) that might not have activated *Cre*-expression. Consequently, a broader view on the Myf5+ population, giving rise to and residing within *Pax7*-expressing cells, could be resolved. Remarkably, this analysis showed that at E11.5 and E14.5, all Pax7+ cells showed a history of *Myf5* expression (Fig. 3E).

Based on the dynamics of expression of these transcription factors, we used *Myf5^Cre^* (Haldar et al., 2007) to delete *Pitx2* before the onset of *Pax7* expression. Immunostaining for Pax7, Pitx2, Myod, Myog and MF20 in *Myf5^Cre^; Pitx2^flox/^* ^Δ^ E14.5 embryos showed no Pax7+ cells, only a few Myod+/Myog+ cells, and no myofibres (Fig. 3F, 3G). Analysis at foetal stages confirmed the lack of EOM in the mutant, albeit a small cluster of Pax7+ cells and myofibres were detected in dorsal sections, at the level of the muscle attachment point to the base of the skull (Fig. S3A-C). As these EOM remnants contained Pitx2+/Pax7+ and Pitx2-/Pax7+ cells and correspond to the site of the initiating EOM anlagen, it is possible that *Myf5^Cre^* was less efficient in this location or they were derived from Mrf4+ cells (Sambasivan et al., 2009).

Given that *Myf5* is expressed briefly in muscle progenitors and downregulated when myoblasts commit to differentiation (Comai and Tajbakhsh, 2014), we used the more downstream commitment marker *Myod* to abolish *Pitx2* expression in myoblasts using *Myod^iCre^* (Kanisicak et al., 2009). Interestingly, myogenesis appeared unperturbed in the EOM in this case (Fig. 4A, 4B). In addition, and in stark contrast to the *Myf5^Cre^* data, similar numbers of *Pax7*-expressing cells were present in controls and *Myod^iCre^;Pitx2^fl/fl^* foetuses (Fig. 4C), despite recombination taking place in >90% of the cells as per *Myod^iCre^;R26^Tomato^* lineage tracing (Fig. S4A, S4B). Taken together, these results show that *Pitx2* is required in *Myf5+* myogenic progenitors for the specification and emergence of the Pax7+ population and is then dispensable upon activation of *Myod*.

**Figure 4.**
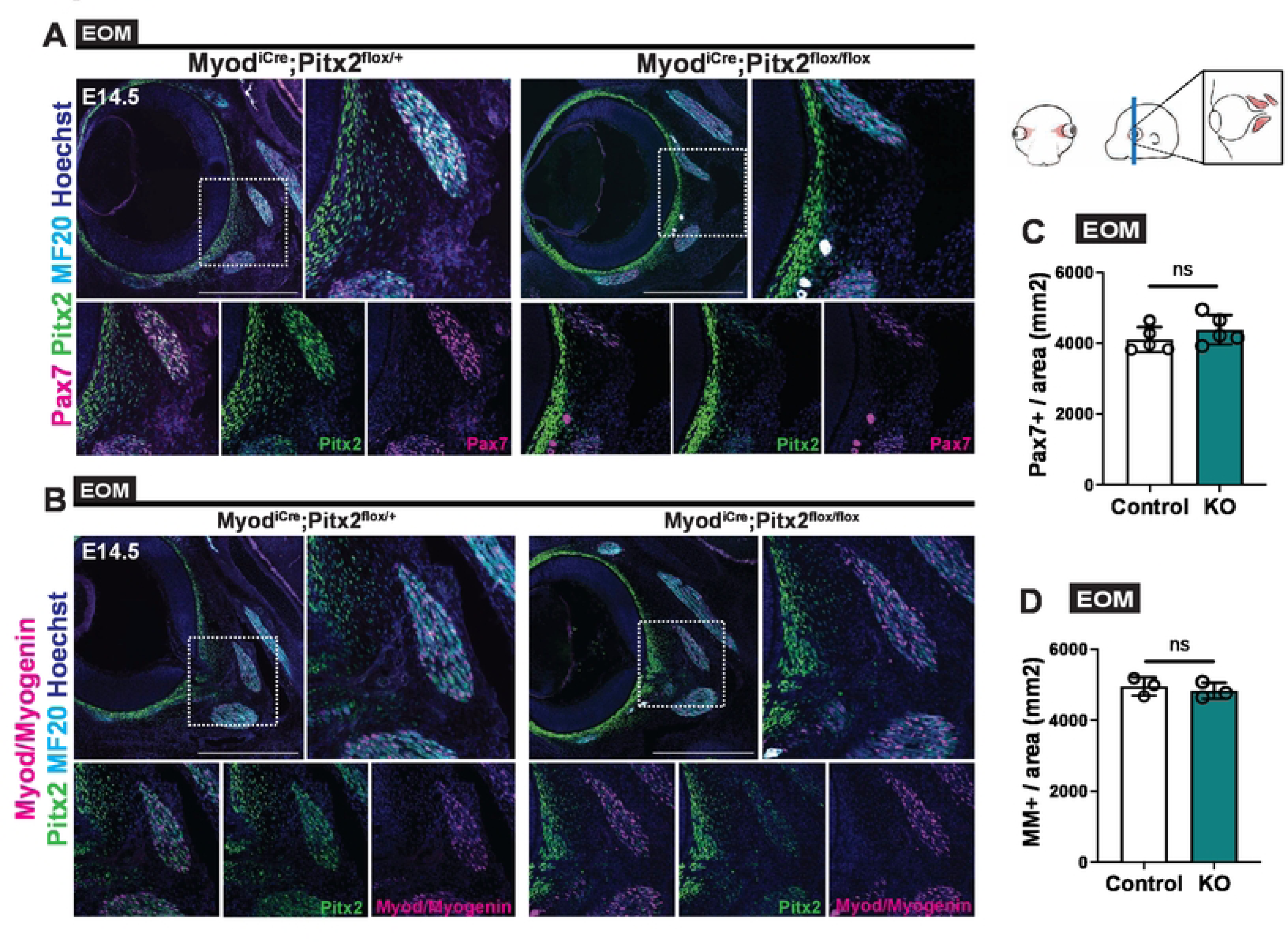
Normal EOM formation following deletion of *Pitx2* in *Myod*-expressing cells. **(A)** Immunostaining of E14.5 EOM sections from *Myod^iCre^;Pitx2^flox/+^* (control) and *Myod^iCre^;Pitx2^flox/flox^* (KO) for Pax7 (magenta), Pitx2 (green) and MF20 (cyan). Bottom panels, higher magnification views of the area delimited with dots. **(B)** Number of Pax7+ cells per area in EOM sections from the immunostaining in (A) (n=4 each). **(C)** Immunostaining of E14.5 EOM sections from *Myod^iCre^;Pitx2^flox/+^* (control) and *Myod^iCre^;Pitx2^flox/flox^* (KO) for Myod/Myogenin (MM) (magenta), Pitx2 (green) and MF20 (cyan). Bottom panels, higher magnification views of the area delimited with dots. **(D)** Number of MM+ cells per area in EOM sections from immunostaining in (C) (n=3). Scale bars: 500μm (A) and (B). Error bars represent SEM. ns, non-significant.

### Temporal requirement for *Pitx2* in EOM stem cells

To assess the role of *Pitx2* specifically in the Pax7+ cells, we generated Tamoxifen-inducible *Pax7^CreERT2^; Pitx2^flox/flox^* mice (conditional KO). Tamoxifen was induced twice at E11.5 and E12.5, at the onset of *Pax7* expression in developing EOM (Horst et al., 2006) (Fig. 3B, 3C), and samples were collected at E17.5 (Fig. 5A). Immunostaining for Pax7, Pitx2 and MF20 showed that although EOM myofibres were present in *Pax7^CreERT2/+^; Pitx2^flox/flox^* foetuses (Fig. 5B), the number of Pax7+ cells was reduced by 31.3% (Fig. 5C), and the proliferating population (Ki67+) was reduced by 16.6% (Fig. 5D) in the conditional KO. No significant perturbations were noted in the HL muscle at this stage. We note that these experiments do not distinguish whether deletion of *Pitx2* impairs the continuous emergence of Pax7+ cells from E11.5, or if the reduced number of Pax7+ cells observed in the foetus is due to reduced proliferation of Pax7+/Pitx2- cells.

**Figure 5.**
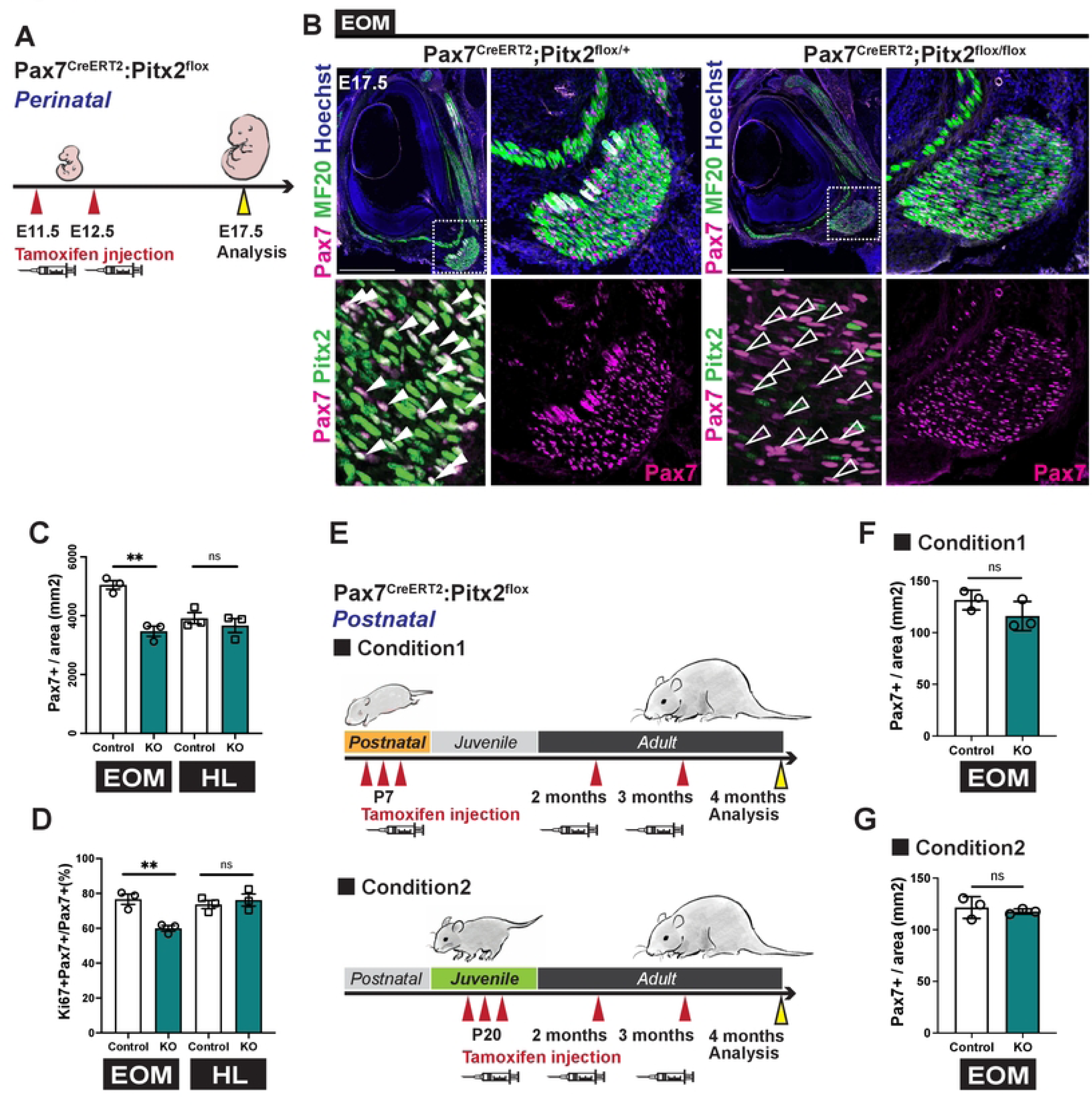
Timing of inactivation of *Pitx2* differently impairs EOM MuSCs. **(A)** Scheme indicating timing of Tamoxifen injections for invalidation of *Pitx2* during embryogenesis. Tamoxifen was injected to *Pitx2^flox/flox^* pregnant females crossed with *Pax7^CreERT2^;Pitx2^flox/+^* male mice and samples collected at E17.5. **(B)** Immunostaining of E17.5 EOM sections from *Pax7^CreERT2^;Pitx2^flox/+^* (control) and *Pax7^CreERT2^;Pitx2^flox/flox^* (cKO) foetuses for Pax7 (magenta), Pitx2 (green) and MF20(cyan). Bottom panels, higher magnification views of the area delimited with dots. White arrowheads indicate Pax7+/Pitx2+ cells and white open arrowheads Pax7+/Pitx2- cells. **(C)** Number of Pax7+ cells per area in EOM and hindlimb (HL) sections from immunostaining in (B) (n=3 each). **(D)** Percentage of Ki67+/Pax7+ cells in total Pax7+ cells in EOM and HL sections (n=3 each). **(E)** Scheme indicating timing of Tamoxifen injections for invalidation of Pitx2 with *Pax7^CreERT2^*. Condition1: Tamoxifen induction from P1-P7. Condition2: Tamoxifen induction from P20. Samples were collected at 4 months of age. **(F)** Number of Pax7+ cells per area in EOM sections from Condition 1 (n=3 each). **(G)** Number of Pax7+ cells per area in EOM sections from Condition 2 (n=3 each). Scale bars: 500μm (B). Error bars represent SEM. ns, non-significant, **P<0.01.

Next, we investigated the role of *Pitx2* in Pax7+ cells postnatally where tamoxifen was injected from the early perinatal (from P1 to P7, condition 1) or weaning (P21, condition 2) stages with additional monthly injections until analysis in the adult (4 months) (Fig. 5E). The rationale was to assure full *Pitx2* deletion from the time of induction, and to avoid a potential rescue by non-deleted cells. In both conditions, there was no difference in the number of EOM Pax7+ cells between the conditional KO and controls (Fig. 5F, 5G), despite an efficient deletion (Fig. S5A-D). Of note, a concomitant deletion of *Pitx2* in myonuclei was observed when Tamoxifen induction was done at P7 (Fig. S5B, S5B’) but not P20 (Fig. S5C, S5C’). This is consistent with our results (Fig. 1C) showing that most of the EOM MuSCs enter quiescence before P20, thus Pax7+ myoblast proliferation and fusion taking place prior to that stage. Therefore, absence of *Pitx2* alone does not result in the loss of Pax7+ MuSCs during homeostasis postnatally.

### Loss of Pitx2 in MuSCs does not account for sparing of EOM in *mdx* mice

High levels of Pitx2 in EOM MuSCs were proposed to be responsible, at least in part, for sparing of EOMs in DMD (Hebert et al., 2013; Verma et al., 2017). Therefore, we generated *mdx^βgeo^; Pax7^CreERT2^; Pitx2^flox/flox^* compound mutant mice (dKO), where *mdx^βgeo^* is a dystrophin null mouse model (Wertz and Füchtbauer, 1998). Tamoxifen was injected 3 times from P1 to P7, followed by monthly injections (from 2 months) until analysis in the adult (Fig. 6A). Immunostaining for Pax7 and Ki67 allowed evaluation of the density and proliferative fraction of Pax7+ MuSCs in EOM and TA (Fig. 6B-F). The number of Pax7+ cells per area was similarly decreased in the EOM of *mdx^βgeo^* and dKO mice compared to controls and cKO (Fig. 6C). Moreover, there were virtually no Ki67+/Pax7+ cells in EOM in *mdx^βgeo^* mice and 3.6% in dKO mice (Fig. 6D), suggesting that EOM MuSCs are not activated in either model at the time points analysed. In contrast, TA muscle from *mdx ^β geo^* mice and dKO showed a significant increase in the number of Pax7+ cells per area, and percentage of proliferative MuSCs (Fig. 6E, 6F) in agreement with the notion that MuSCs undergo activation in *mdx* mice during cycles of muscle degeneration and regeneration (Georgieva et al., 2022).

**Figure 6.**
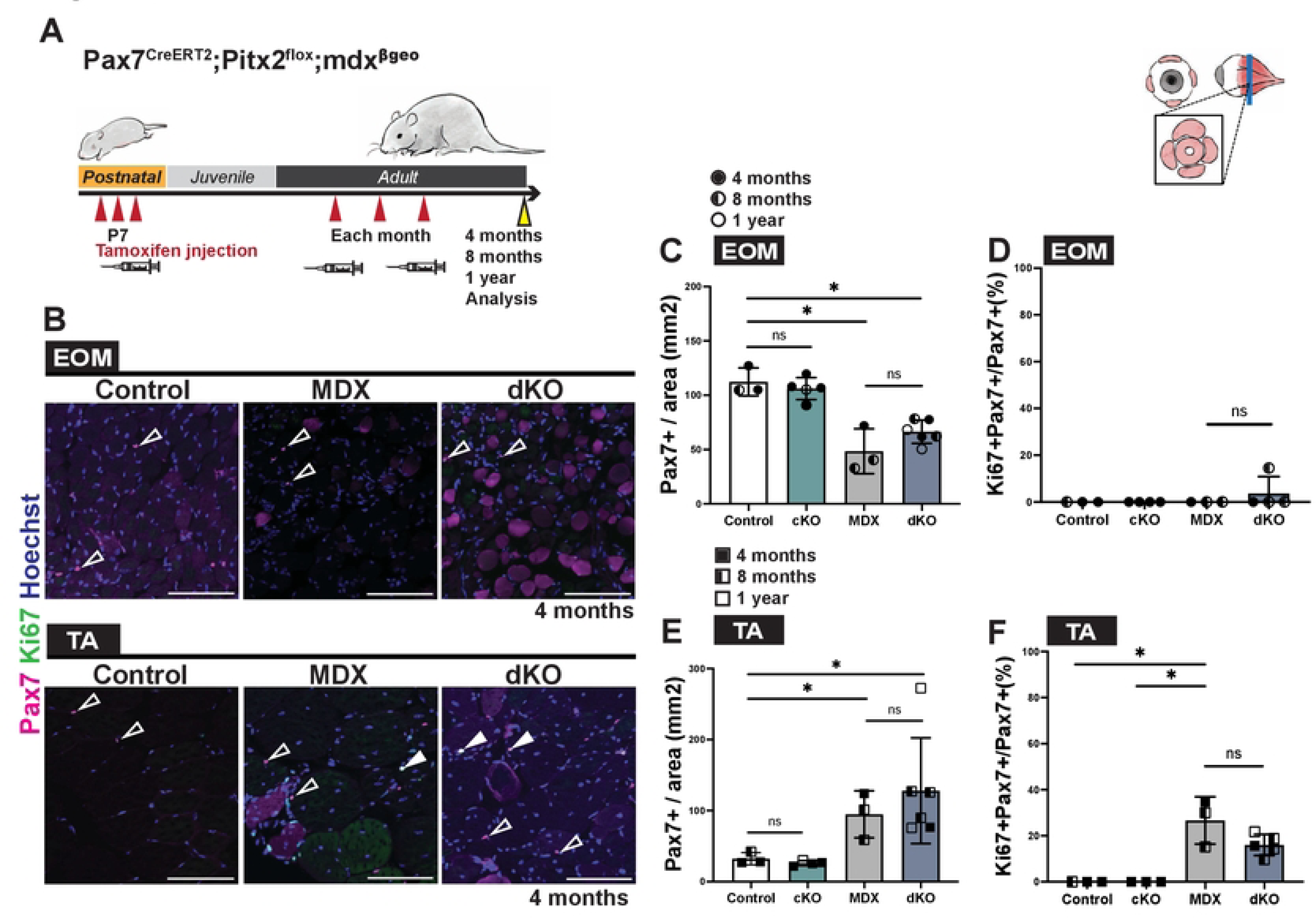
Absence of proliferation in EOM of *mdx* and *mdx;Pitx2* KO mice. **(A)** Scheme indicating timing of Tamoxifen injections on *Pax7^CreERT2^;Pitx2^flox^*; *mdx^βgeo^* and control mice at postnatal and adult stages. Samples were collected at 4 months, 8 months and 1 year for analysis. **(B)** Immunostaining of EOM and TA sections from 4 months old Control (*Pitx2^flox/flox^* or *Pitx2^flox/+^*), cKO, *mdx^βgeo^* (MDX) and *Pax7^CreERT2^;Pitx2^flox/flox^; mdx* (dKO) mice for Pax7 (magenta) and Ki67 (green). White arrowheads point to Pax7+/Ki67+ cells; open arrowheads to Pax7+/Ki67-cells. **(C)** Number of Pax7+ cells per area on EOM sections from immunostaining in (B) (Control, and MDX, n=3; cKO, n=4; dKO, n=6). **(D)** Percentage of Ki67+/Pax7+ cells over total Pax7+cells on EOM sections from immunostaining in (B) (Control, and MDX, n=3; cKO, n=4; dKO, n=4). **(E)** Number of Pax7+ cells per area on TA sections from immunostaining in (B) (Control, and MDX, n=3; cKO, n=4; dKO, n=6). **(F)** Percentage of Ki67+/Pax7+ cells over Pax7+cells on TA sections from immunostaining in (B) (Control, cKO and MDX, n=3; dKO, n=5). 4 months old (Control, MDX and dKO) and 1 year old (dKO and cKO) animals were littermates. 8-month-old mice, dKO and control mice were littermates while MDX were generated in independent litters. All mice were treated with Tamoxifen. Scale bars: 100μm (B) Error bars represent SEM. ns, non-significant, *P<0.05.

Constitutive deletion of *Pitx2* in myofibres of *mdx4cv* mice was reported to result in severe dystrophic changes in EOM compared to a mild fibre phenotype observed in the limb (Verma et al., 2017). Thus, we assessed the number of centrally nucleated myofibres in the EOM of both *mdx^βgeo^* and dKO mice (Fig. S6A, S6B). Although in the dKO a large fraction of myonuclei is Pitx2- as they were incorporated from Pitx2-deleted Pax7+ cells after P7 (Fig. S5B), only sparse centrally located myonuclei were observed (2.0% in dKO compared to 0.6% in *mdx ^β geo^* Fig. S6A, S6B). This is at odds with the high numbers of centrally nucleated fibres normally observed in *mdx^βgeo^* TA muscles (Fig. S6A), which can reach up to ∼60% (Lau et al., 2018). Next, we calculated the cross-sectional area of EOM myofibres in the 4 recti muscles from WT, *mdx^βgeo^*, *Pitx2* cKO and dKO and classified them by size (Fig. S6C). While a trend towards larger myofibre size was observed in the EOM of *mdx ^β geo^* mice, dKO mice showed some heterogeneity with a slight shift towards smaller and larger size fibres (Fig. S6C). Altogether, *Pitx2* appears to play a minor role in EOM MuSCs in a dystrophic background.

## Discussion

Pax7 plays a critical role in the maintenance of mouse muscle stem cells from late fetal stages and it is the most widely used marker for adult MuSCs and their ancestors from mid-embryogenesis (Gunther et al., 2013; Kassar-Duchossoy et al., 2005; Lepper et al., 2009; Relaix et al., 2006; Seale et al., 2000; von Maltzahn et al., 2013). Pax7 is required for maintenance and function of the MuSC pool from late fetal stages in mouse (Gunther et al., 2013; Murphy et al., 2011; von Maltzahn et al., 2013). Adult MuSCs in the head also express Pax7, yet they also retain the expression of the early mesodermal markers characteristic of each location (Evano et al., 2020; Harel et al., 2009; Ono et al., 2010; Sambasivan et al., 2009). In this study, we investigated the functional relationships between *Pax7*, *Pitx2* and *Myf5* in the emergence and maintenance of MuSCs. Our data reveal different temporal requirements for *Pitx2* during prenatal and postnatal stages, and during lineage progression, for the emergence of the EOM Pax7+ stem cells.

### Interplay between Pitx2, Myf5 and Pax7 in emergence of EOM stem cells

In the trunk, both at the dermomyotome and limb levels, expression of *Pax3* and *Myf5* precede the expression of *Pax7* (Comai and Tajbakhsh, 2014; Jostes et al., 1990; Kassar-Duchossoy et al., 2005; Tajbakhsh et al., 1997), with *Pax3* expression being then rapidly downregulated during myogenic cell commitment (Myod+ cell). Following the establishment of a muscle anlagen (uncommitted and differentiated cells), *Pax7* is expressed from mid-embryogenesis in the upstream undifferentiated population, and it persists to give rise to adult MuSCs (Kassar-Duchossoy et al., 2005; Relaix et al., 2005). In the head of vertebrate models analysed, Pax7+ myogenic cells were reported to emerge de novo, as *Pax3* is not expressed in head MuSCs (Horst et al., 2006; Nogueira et al., 2015). Instead, and like the role of *Pax3* in the trunk and limbs, *Pitx2* is critically required cell-autonomously to maintain myogenic progenitors in an immature state, ensure their survival, and to activate the MRFs (Zacharias et al., 2011). Differently to the role of *Pax3* in the trunk, *Pitx2* expression persists following activation of the MRFs and myogenic commitment in the developing EOMs, and it is required for myogenic cell survival until the end of the embryonic period (Sambasivan et al., 2009). The expression of Pitx2 also in adjacent mesenchymal cells (Comai et al., 2020; Gage et al., 1999) has complicated the analysis and interpretation of the cell-autonomous roles of this gene in EOM development. In addition, conditional inactivation of *Pitx2* in neural-crest-derived cells does not affect the specification of the EOM primordia, but it results in severe muscle patterning defects (Evans and Gage, 2005).

We demonstrated herein a temporal requirement for *Pitx2* during lineage progression: deletion of this gene in muscle progenitors (*Myf5^Cre^*) results in the absence of myofibres and upstream cells in the embryo, while deletion in myoblasts (*Myod^iCre^*) does not. Further, deletion of this gene in Pax7+ cells led to a reduction in the number of EOM MuSCs in the foetus, whereas *Pitx2* function appears to be dispensable for EOM MuSC maintenance in the adult. In this muscle group, Myf5 and Mrf4 also play critical roles as EOMs and the upstream population are absent in *Myf5;Mrf4* double mutants (Sambasivan et al., 2009; Tajbakhsh and Buckingham, 2000). Collectively, these observations indicate that EOM specification and differentiation has evolved a unique gene regulatory network where EOM progenitor survival as well as activation of the downstream myogenic factors are dependent on either *Myf5* or *Mrf4*, and that *Pitx2* alone cannot override their role. Therefore, it is possible that the *Pitx2* survival function is progressively relayed onto *Myf5* and/or *Pax7*. A double *Pax7;Pitx2* KO in the MuSC lineage would be required to formally address this hypothesis. Moreover, while Pitx2 binds directly to the promoters of *Myf5* and *Myod* (Zacharias et al., 2011), it is unclear whether it can directly activate *Pax7*, or if other TFs are necessary. Of note, sc-RNAseq data from our lab (Grimaldi et al., 2022) revealed several novel TFs that are specifically expressed in EOM progenitors. Whether those factors are required for the emergence of the myogenic population in the EOM requires further investigation. Moreover, a recent study showed that *Six1* and *Six2* genes are required for craniofacial myogenesis by controlling the engagement of unsegmented cranial paraxial mesodermal cells in the myogenic pathway (Wurmser et al., 2023). It remains unclear if expression of these genes in adult MuSCs would be required for *Pax7* maintenance in the EOM.

### Adult EOM MuSCs do not proliferate during homeostasis despite coexpression of Pax7 and Pitx2

Activated EOM MuSCs have greater proliferative and self-renewal abilities *in vitro* compared to those in the limb (Di Girolamo et al., 2023; Stuelsatz et al., 2015). Moreover, expression of *Pitx2* in postnatal EOM myogenic cells (CD34+/Sca1−/CD31−/CD45− lineage), at higher levels than limb MuSCs, was reported to contribute to the proliferative and stress resistance properties of EOM MuSCs *in vitro* (Hebert et al., 2013; Kallestad et al., 2011). However, with respect to the proliferative status of EOM MuSCs *in vivo*, we note several discrepancies. In some species, EOM MuSCs were suggested to chronically proliferate *in vivo* (Kallestad et al., 2011; McLoon and Wirtschafter, 2002, 2003), but these studies generally lacked co-immunostainings for Pax7 and proliferation markers. In addition, EOM MuSCs appeared to contribute to new myofibre myonuclei in homeostasis at higher frequency than that observed for limb muscles (Keefe et al., 2015; McLoon and Wirtschafter, 2002; Pawlikowski et al., 2015). In this study, we performed co-immunostaining for Pax7 and Ki67 and found that EOM MuSCs entered quiescence earlier than those in TA muscle, and we did not observe proliferative EOM MuSCs during adult homeostasis. These observations agree with a previous study (Stuelsatz et al., 2015) showing that freshly isolated mouse EOM MuSCs are not proliferative. While the origin of the discordance with lineage tracing studies is unclear, one possibility could be that EOM MuSCs contribute to myofibres without cell cycle entry and/or the genetic constructs used as *Cre* drivers are unexpectedly expressed in EOM myofibres.

It was also proposed that Pitx2+ myogenic cells do not co-express *Pax7* but express *Myod* in human EOM, and these cells could then represent a second progenitor population involved in EOM remodeling, repair, and regeneration (Verma et al., 2017). Given our observation that more than 92.5% of Pax7+ MuSCs coexpress *Pitx2*, this might highlight species specific differences or antibody sensitivity and tissue processing issues in the different studies.

### Deletion of Pax7 results in loss of the EOM MuSC population

Several studies analysed the limb and trunk phenotypes of constitutive (Kuang et al., 2006; Oustanina et al., 2004; Relaix et al., 2006; Seale et al., 2000) and inducible (Gunther et al., 2013; Lepper et al., 2009; von Maltzahn et al., 2013) *Pax7* mutants. While MuSC progenitors are initially specified in homeostatic conditions (Oustanina et al., 2004), the progressive loss of MuSCs in newborn mice appears to be due to proliferation defects of MuSCs and cell death (Oustanina et al., 2004; Relaix et al., 2006). Elimination of *Pax7* in adult MuSCs results in MuSC loss likely due to enhanced differentiation and reduced heterochromatin condensation in the remaining cells (Gunther et al., 2013) and severe deficits in muscle regeneration (Gunther et al., 2013; Kuang et al., 2006; Oustanina et al., 2004; von Maltzahn et al., 2013). The phenotypes of Pax7 null cells during in vitro activation were somewhat more divergent, depending on the system used (isolated myoblasts, myofibers, clonogenic assays) including absence or reduction in the number of myoblasts derived from Pax7KO cells together, increased or reduced differentiation and/or commitment to alternative fates (Kuang et al., 2006; Oustanina et al., 2004; Relaix et al., 2006; Seale et al., 2000; von Maltzahn et al., 2013).

As EOMs are not amenable to in vivo muscle injury, in this study we examined the *in vivo* phenotype of *Pax7^nGFP/nGFP^* null EOM compared to heterozygous controls at E14.5 and P21. The perdurance of the GFP allowed us to trace the progenitors and more differentiated cells. We highlight several differences between the EOM and TA muscles. First, in P21 control samples, EOM displayed a higher number of Pax7+ cells per area, but not per fibre compared to TA. This agrees with previous data showing higher number of MuSCs per volume unit in the EOM (Verma et al., 2017), which might be related to their smaller fibre size. Second, a greater reduction of GFP*+* cells per area was noted in *Pax7* null EOM compared to TA, suggestive of a more relevant role for *Pax7* in the maintenance of MuSCs without a history of *Pax3* expression. Notably, no differences in the number of GFP+ cells per area neither in the proportion of Pitx2+GFP+ cells were observed in the fetal period between control and mutant EOMs, suggesting that other factors different than Pax3 and Pitx2 may ensure the specification of this cell population. Previous studies showed that Pax7 is required for in limb MuSC precursors to express genes normally associated with functional MuSCs such as Syndecan4 and CD34 (Kuang et al., 2006; Oustanina et al., 2004). Here, we further show that expression of CalcR, another MuSC marker (Fukada et al., 2007), was fully impaired in EOM and TA from Pax7 null. In addition, while the fraction of committed GFP*+* cells in Pax7null EOM display a marked increase compared to controls, no significant changes were observed in mutant and control TA at P21. This observation probably reflects differences on the pace of lineage progression between these muscles in vivo. It would be of interest in future studies to assess whether the downstream targets of *Pax7* in MuSCs of cranial and somite origin are similar and to identify complementary TFs that confer these distinct MuSC phenotypes.

### Absence of proliferation in EOM of *mdx* and *mdx;Pitx2* KO mice

Dystrophin protein is expressed in differentiated myofibres where it is required for sarcolemmal integrity, but it is also expressed in activated MuSCs, where it regulates MuSC polarity and the mode of cell divisions (Dumont et al., 2015; Wang et al., 2019; Weng et al., 2019). Moreover, several mechanisms of MuSC dysfunction have been described in DMD (Filippelli and Chang, 2022), including severe proliferation defects and premature senescence of chronically activated MuSCs (Blau et al., 1983; Chang et al., 2021; Sugihara et al., 2020; Taglietti et al., 2023). Notably, in a preclinical rat model of DMD, expression of thyroid-stimulating hormone receptor (Tshr) was shown to protect EOM MuSCs from entering senescence and for them to remain in a proliferative state (Taglietti et al., 2023).

In this study, we found that the number of Pax7+ cells/area were reduced in EOM of *mdx* and *mdx;Pitx2* KO mice, and that these cells were not proliferative. The *mdx* mice have a relatively normal lifespan and have a milder phenotype compared to rats, dogs, and human (McGreevy et al., 2015; Taglietti et al., 2022). Thus, the absence of proliferation of *mdx* EOM MuSCs might reflect a complete sparing of these muscles in the murine model. Alternatively, it might reflect activation and fusion of the MuSC population without prior entry in the cell cycle given that the number of Pax7+ cells/area were reduced in *mdx* and *mdx;Pitx2* mice compared to control EOMs. Moreover, we observed similar reductions in the number of MuSCs in the EOM of *mdx* and *mdx;Pitx2* mice. As the *Pax7^CreERT2^;Pitx2^fl/fl^* EOMs were unaffected when *Pitx2* was deleted postnatally, these data suggest that the reduction in the number of EOM MuSCs is *mdx* but not *Pitx2* dependent.

Finally, some insights into the potential role of *Pitx2* in the sparing of EOMs in *Dmd* come from a preliminary study showing that in the absence of *Pitx2* expression in myofibres of *mdx4cv* mice, the EOMs succumb to dystrophic changes that are even more severe than those seen in the limb muscles of the same mice (Verma et al., 2017). In the present study, we deleted *Pitx2* in MuSCs at an early postnatal stage (<P7), where considerable fusion is ongoing and thus MuSC-derived myoblasts actively contribute to new myonuclei. While this resulted in a small percentage of centrally nucleated fibres and a more heterogeneous distribution of fibre sizes (shift towards smaller and larger fibres), the resulting phenotype was less severe than when *Pitx2* was deleted with a constitutive myofibre *Cre* driver (Verma et al., 2017). However, the latter study was performed with 18 months old mice, therefore a long-term phenotype upon deletion of *Pitx2* in MuSCs cannot be excluded. Performing deletion of *Pitx2* prenatally in MuSCs or timed deletions within myofibres using a conditional fibre *Cre* driver mouse would help elucidate the relative and roles of *Pitx2* in each compartment.

In summary, how some muscles are spared in myopathies, such as the EOM in dystrophic mice, remains unresolved and under debate. Our studies on the prenatal and postnatal development of susceptible and spared muscles provides insights into this process regarding the roles of key transcription factors that specify and maintain the muscle stem cell populations in these muscles.

## MATERIALS AND METHODS

### Mouse strains

Animals were handled as per European Community guidelines and the ethics committee of the Institut Pasteur (CETEA) approved protocols (APAFIS#41051-202302204207082). The following strains were previously described: *mdx-βgeo* (Wertz and Füchtbauer, 1998), *Myf5^Cre^* (Haldar et al., 2007), *Myf5^nlacZ^* (Tajbakhsh et al., 1996), *Pax7^CreERT2^* (Murphy et al., 2011), *Myod^iCre^* (Kanisicak et al., 2009), *mdx^β geo^* (Wertz and Füchtbauer, 1998), *Pax7^nGFP^* (Sambasivan et al., 2009) and *Pitx2^flox^* (Gage et al., 1999), in which the DNA binding homedomain region common to Pitx2a/b/c isoforms is flanked with LoxP sites. Mice were kept on a mixed genetic background (B6D2F1, Janvier Labs). Mouse embryos and foetuses were collected at embryonic day (E) E14.5 and E17.5, with noon on the day of the vaginal plug considered as E0.5. To generate conditional KOs, *Pax7^CreERT2/CreERT2^ ; Pitx2^flox/+^*, *Myf5^Cre/+^ ; Pitx2^flox/+^* and *Myod^iCre/+^ ; Pitx2^flox/+^* males were crossed with *Pitx2^flox/flox^* or *Pitx2^flox/+^* females. The cross with *Myf5^Cre/+^ ; Pitx2^flox/+^* male and *Pitx2^flox/flox^* or *Pitx2^flox/+^* females generated *Myf5^Cre/+^ ; Pitx2^flox/^* ^Δ^ fetuses as conditional KOs. To generate dKO (*Pax7^CreERT2/+^ ; Pitx2^flox/+^; mdx^βgeo^*), *Pax7^CreERT2/CreERT2^ ; Pitx2^flox/+^* male was crossed with *Pitx2^flox/flox^ ; mdx^βgeo/+^* or *Pitx2^flox/+^; mdx^βgeo/+^* females.

### Immunofluorescence

Embryonic tissues for cryosections were fixed for 2h in 4% paraformaldehyde (PFA) in PBS (Electron Microscopy Sciences, Cat #:15710) with 0.5% Triton X-100 at 4°C. Adult muscles for tissue sections were fixed for 2h in 1% PFA in PBS with 0.1% Triton X-100 at 4°C. After PBS washes for a few hours, samples were equilibrated in 30% sucrose in PBS overnight at 4°C then embedded in OCT compound (Sakura Fineteck, Cat #:4583). Cryosections (16μm) were permeabilized with 0.5% Triton in PBS for 5 min at RT and blocked for 1h at RT in Blocking solution (10% Goat serum, 1% BSA and 0.5% Triton X-100 in PBS). Primary antibodies used are: chicken polyclonal anti-GFP (Abcam, Cat#: ab13970, dilution 1:1000), mouse monoclonal anti-Myod (BD-Biosciences, Cat#: 554130, dilution 1:500), mouse monoclonal anti-Pax7 (DSHB, Cat# Pax7, dilution 1:20), mouse monoclonal anti-Myog (DSHB, Cat# F5D, dilution 1:20), rabbit polyclonal anti-Pitx2 (Abcam, Cat# ab221142, dilution 1:750), rabbit polyclonal Anti-Ki67 (Abcam, Cat# ab16667, dilution 1:500), mouse monoclonal anti-MF20 (DSHB, Cat# MF 20, dilution 1:20) rabbit polyclonal anti-Calcitonin Receptor (Bio-RAD, Cat# AHP635, dilution 1:2000), chick anti-GFP (abcam, Cat# 13970, dilution 1:1000), rabbit β-gal antibody (MP 559761, dilution 1/500), rabbit anti DsRed (clontech 632496, dilution1:200) for tdTomato detection and rabbit polyclonal anti-Laminin (Sigma-Aldrich, Cat# L9393, dilution 1:1000). Following washes in PBS-T (0.05% Tween20 in PBS), fluorescent secondary antibodies (dilution 1:500) and Hoechst (Thermo Scientific, Cat. #:H3570, dilution 1:10000) were diluted in Blocking solution and incubated for 1h at RT. Images were acquired by Spinning disk Ti2E (Nikon). Whole Mount immunostaining and clearing were performed as described (Comai et al., 2020). Images were acquired with a confocal microscope (LSM700) and 3D reconstructions performed on Imaris.

### Tamoxifen injection

Tamoxifen injections were done to induce *Cre^ERT2^*-mediated recombination in *Pax7^CreERT2^;Pitx2^flox/flox^* mice. A solution of 15-20mg/ml Tamoxifen (Sigma-Aldrich, Cat# T5648) in 5% ethanol in sunflower oil was prepared by vortexing and rolling at 4°C in the dark and kept for up to one week at 4°C. For the induction at embryonic stages, pregnant female mice were treated with 150μl tamoxifen solution (20 mg/ml) by gavage at the indicated timepoints. For induction of before P7, pups were injected subcutaneously with 10μl tamoxifen solution (15mg/ml). For induction of at P20, pups were injected intraperitoneally with 50μl tamoxifen solution (20mg/ml). Adult mice were treated with 150μl tamoxifen solution (20 mg/ml) by gavage monthly from 2 months until the time of analysis. Tamoxifen was injected in all animals regardless of their genotypes (Control, MDX, cKO and dKO).

### Image analysis and statistics

Quantifications were performed using Fiji (https://imagej.net/software/fiji/). Barplots were generated using Prism (https://www.graphpad.com/features). Fibre sizes and numbers were measured automatically by Cellpose (https://cellpose.readthedocs.io/en/latest/) and Fiji. All data are presented as the mean±s.d. with n (number of animals analysed) ≥3. Statistical significance was assessed using Student’s t-test.

## Acknowledgements

We acknowledge funding support from the Institut Pasteur, Agence Nationale de la Recerche (Laboratoire d’Exellence Revive, Investissement d’Avenir; ANR-10-LABX-73 to ST and ANR-21-CE13-0005 MUSE to GC), Association Française contre les Myopathies (Grant #20510 to ST and #23201 to GC), and the Centre National de la Recherche Scientifique. M.K. was supported by a Post-Doctoral Fellowship from Uehara Memorial Foundation (Grant# 202231020). We gratefully acknowledge the UtechS Photonic BioImaging (Imagopole), C2RT, Institut Pasteur, supported by the French National Research Agency (France BioImaging; ANR-10–INSB–04; Investments for the Future) for support in conducting this study. We thank Dounia Bouragba and Sayna Miri for initial contributions to this study.

## Author contributions

GC and ST conceived the study. MK and GC performed experiments and analysed data. GC, MK, and ST wrote the manuscript. GC and ST provided funding.

## SUPPLEMENTARY FIGURE LEGENDS

**Supplementary Figure 1.**
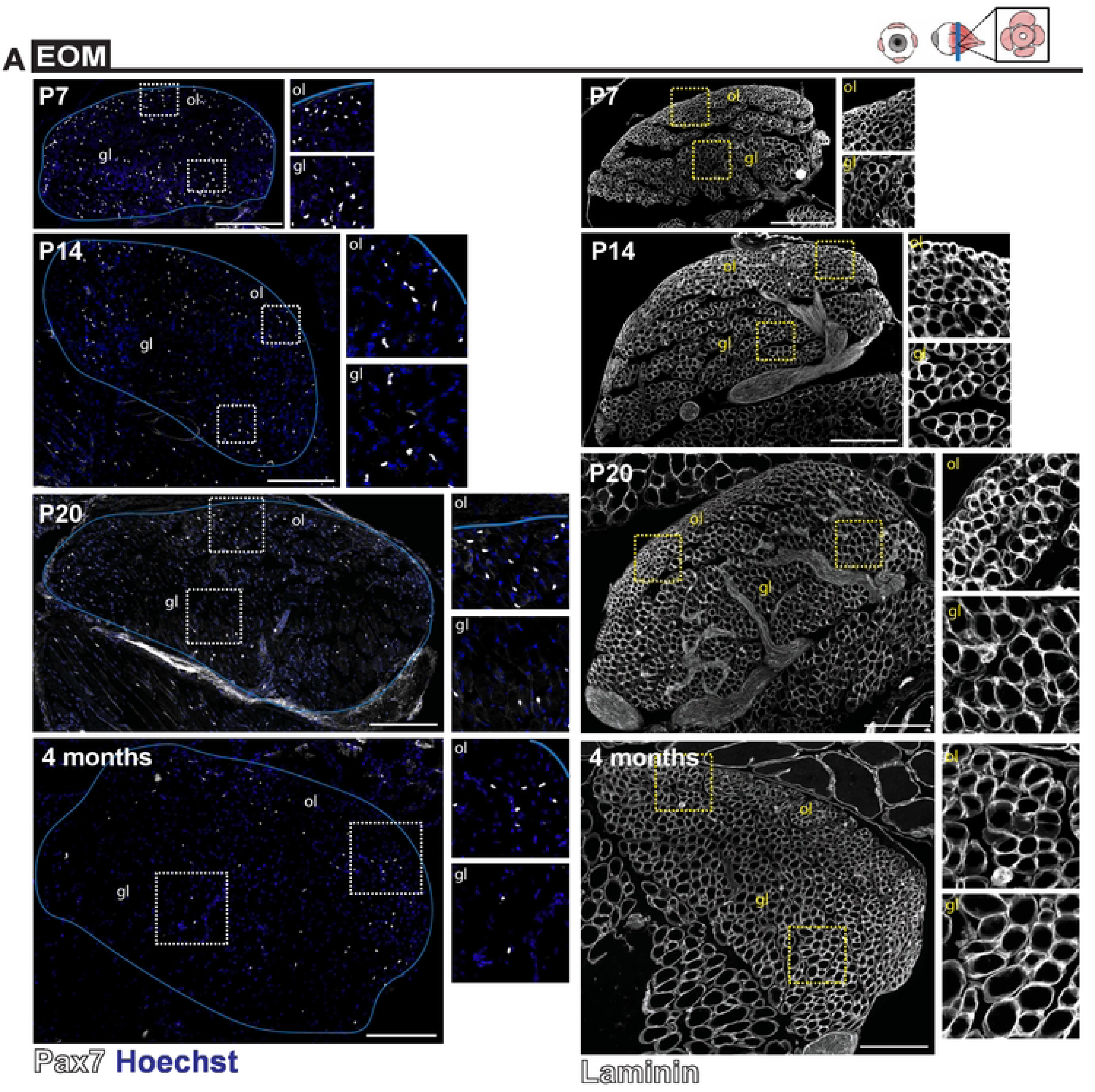
Analysis of orbital and global EOM layers postnatally. **(A, B)** Immunostaining of P7, P14, P20 and 4 months old mice EOM sections for (A) Pax7 (white) and Hoechst nuclei staining or (B) Laminin (white). Global layer (gl) and orbital layer (ol). Higher magnification views delimited with dots. Scale bars: 200μm

**Supplementary Figure 2.**
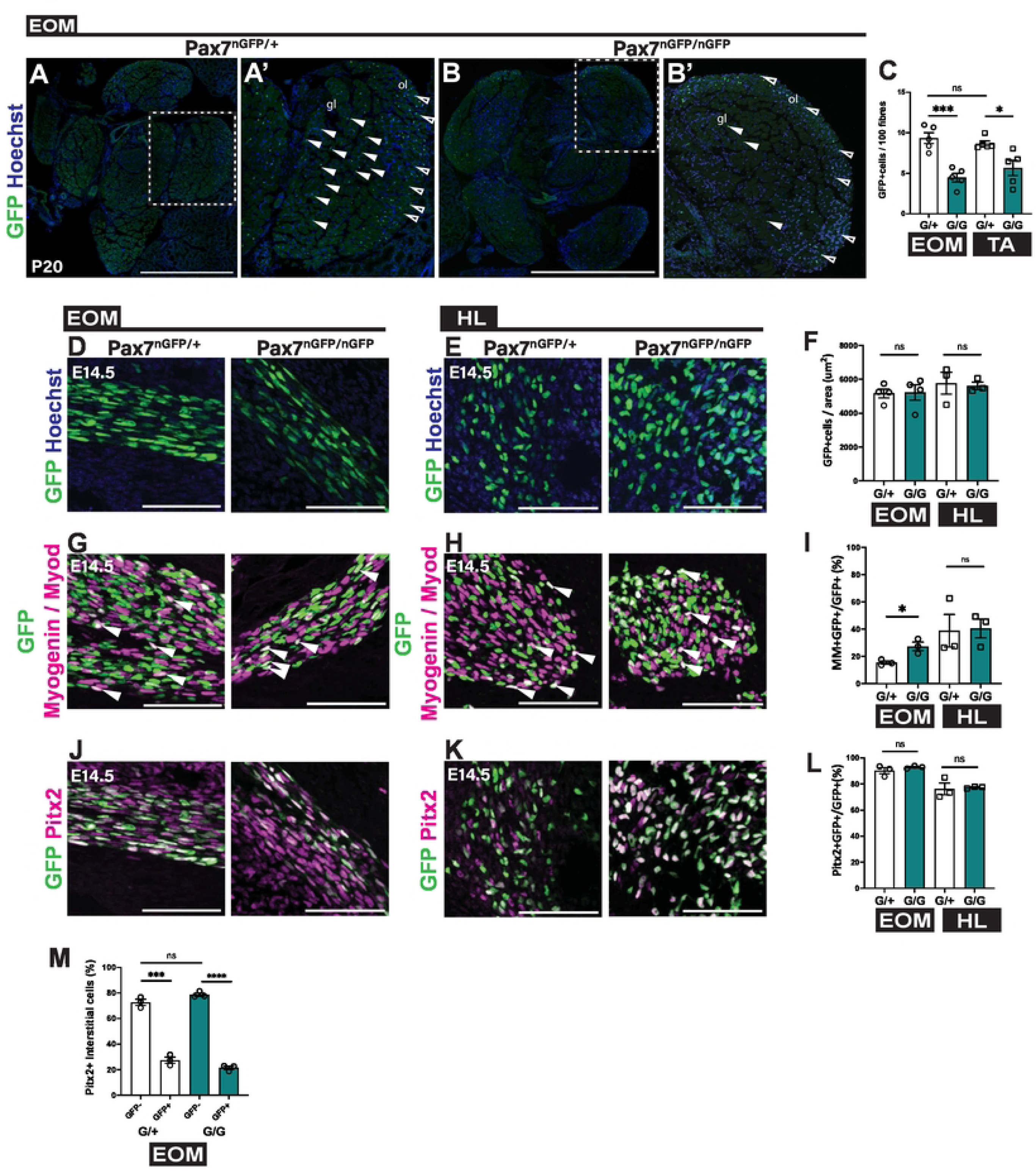
Role of *Pax7* in the maintenance of EOM myogenic cells. **(A, B)** Immunostaining of EOM sections from *Pax7^nGFP/+^* and *Pax7^nGFP/nGFP^* mice at P20 for GFP (green), and Hoechst (blue). **(A’, B’)** Higher magnification views of the area delimited with dots. White arrowheads indicate GFP+ cells in global layer (gl); open arrowheads indicate GFP-cells in orbital layer (ol). **(C)** Number of GFP+ cells per 100 fibres of EOM and TA sections at P20 (n=5 each). **(D,E)** Immunostaining of EOM and HL (hindlimb) sections from *Pax7^nGFP/+^* and *Pax7^nGFP/nGFP^* mice at E 14.5 for GFP (green). (F) Number of GFP+ cells per area in EOM and HL sections from immunostaining in (D,E). (EOM n=4, TA n=3). **(G,H)** Immunostaining of EOM and HL (hindlimb) sections from *Pax7^nGFP/+^* and *Pax7^nGFP/nGFP^* mice at E 14.5 for GFP (green) and Myod/Myogenin (MM) (magenta). White arrowheads indicate MM+/GFP+ cells. **(I)** Percentage of MM+/GFP+ cells over total GFP+ cells in EOM and HL sections from immunostaining in (G,H) (n=3 each). **(J,K)** Immunostaining of EOM and HL (hindlimb) sections from *Pax7^nGFP/+^* and *Pax7^nGFP/nGFP^* mice at E 14.5 for GFP (green) and Pitx2 (magenta), **(L)** Percentage of Pitx2+/GFP+ cells over total GFP+ cells in EOM and HL sections from immunostaining in (J,K) (n=3 each). **(M)** Percentage of GFP+/Pitx2+ interstitial cells and GFP-/Pitx2 interstitial cells in EOM sections at P20 (n=3 each). Error bars represent SEM. ns, non-significant, *P<0.05, **P<0.01, ***P<0.001, ****P<0.0001.

**Supplementary Figure 3.**
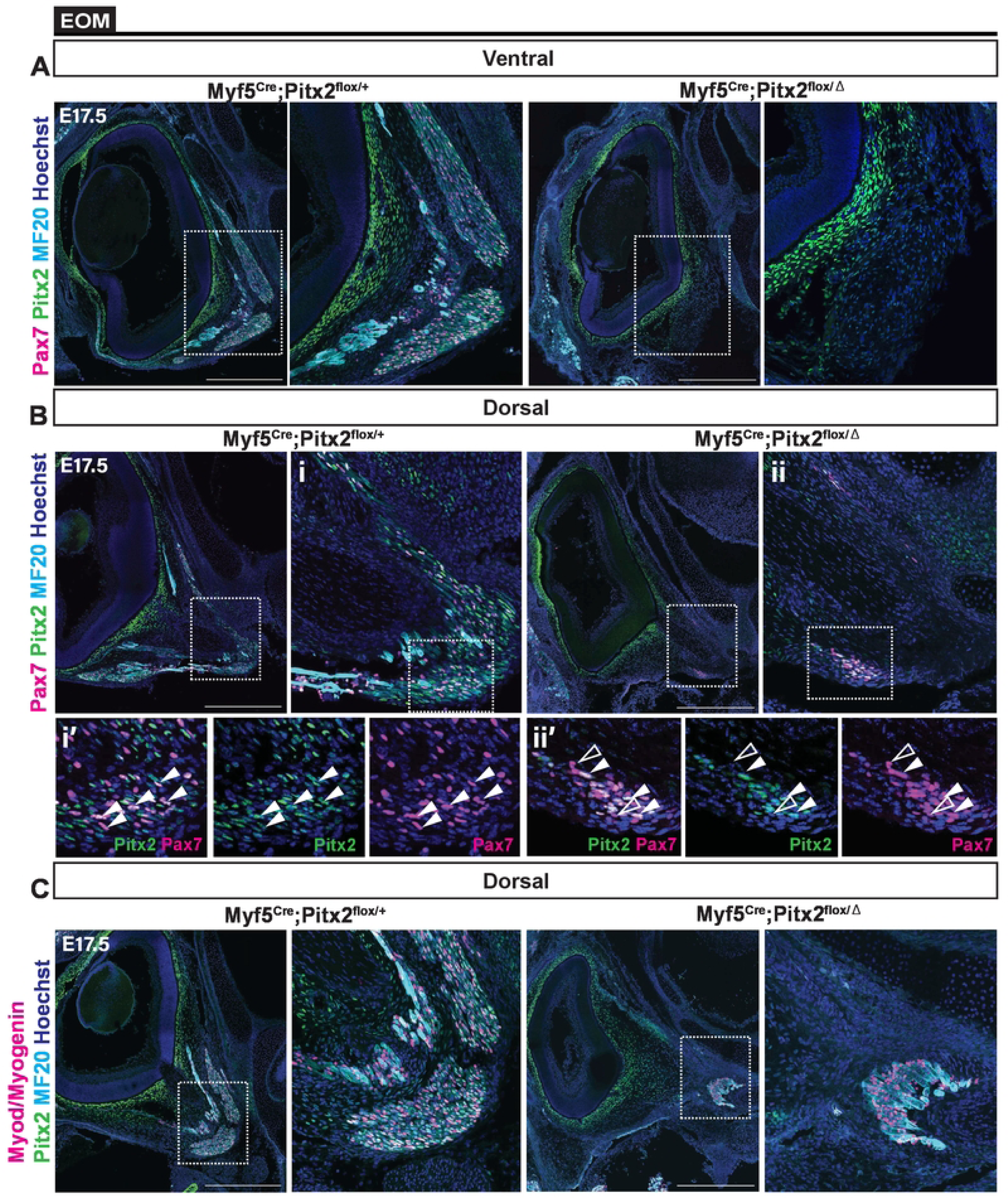
Loss of EOM in *Myf5^Cre^; Pitx2^flox/flox^* mice. **(A,B)** Immunostaining of EOM sections from *Myf5^Cre^;Pitx2^flox/+^* (control) and *Myf5^Cre^;Pitx2^flox/^* ^Δ^ (KO) at E17.5 for Pax7 (magenta), Pitx2 (green) and MF20 (cyan). White arrow heads indicate Pax7+/Pitx2+ cells; open arrow heads indicate Pax7+/Pitx2- cells. Right panels, higher magnification views of the area delimited with dots. **(C)** Immunostaining of EOM sections from *Myf5^Cre^;Pitx2^flox/+^* (control) and *Myf5^Cre^;Pitx2^flox/^* ^Δ^ (KO) at E17.5 for Myod/Myogenin (magenta), Pitx2 (green) and MF20 (cyan). Bottom panels, higher magnification views of the area delimited with dots. Samples were evaluated in ventral (A) and dorsal (B,C) anatomical locations. Scale bars: 500μm (A-C).

**Supplementary Figure 4.**
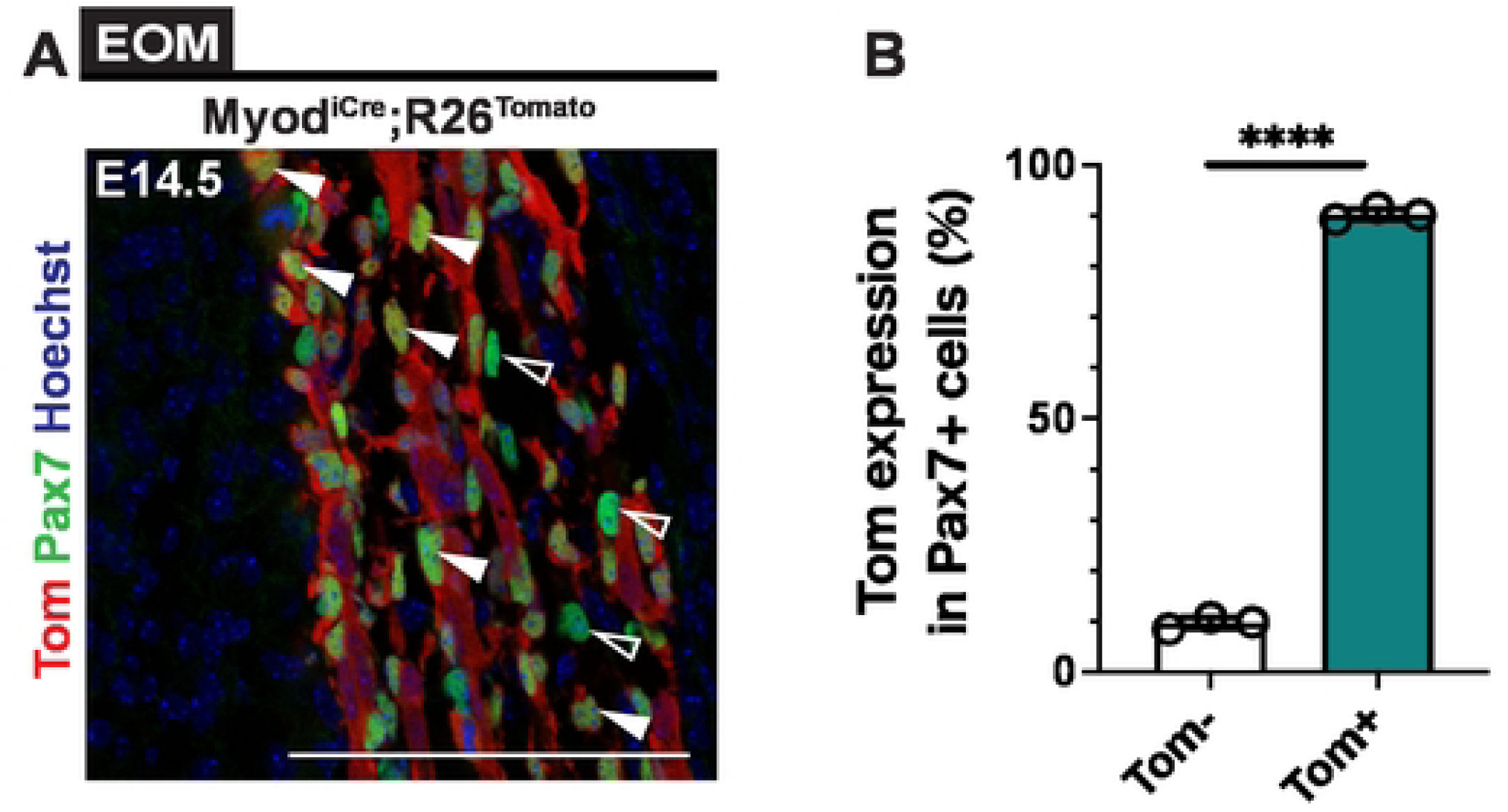
Recombination frequency of cells from *Myod^iCre^;R26^Tomato^* lineage tracing in EOM. **(A)** Immunostaining of EOM sections from *Myod^iCre^;R26^Tomato^* at E14.5 for Pax7 (green), Tomato (red), Pitx2 (green) with Hoechst (blue). White arrowheads indicate Tomato+/Pax7+ cells; open arrowheads indicate Tomato-/Pax7+ cells. **(B)** Percentage of Tomato+ and negative Pax7+ cells in EOM from immunostaining in (A) (n=3 each). Scale bars: 100μm (A). Error bars represent SEM. ****P<0.0001.

**Supplementary Figure 5.**
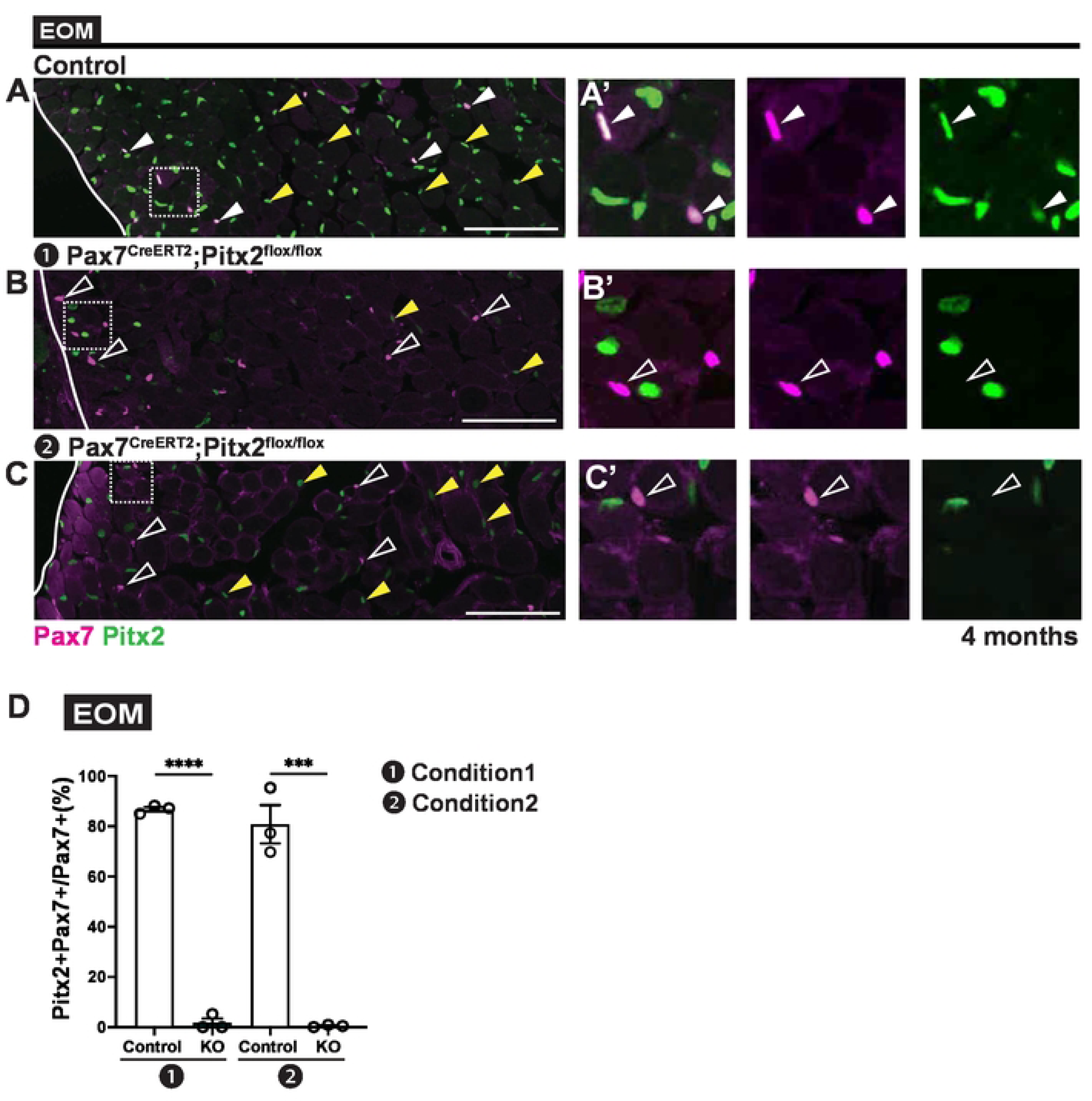
Evaluation of *Pitx2* deletion efficiency with *Pax7^CreERT2^* mice. **(A-C)** Immunostaining of EOM sections from 4 months old Control (*Pitx2^flox^*^/+^) and *Pax7^CreERT2^;Pitx2^flox/flox^* (KO) mice from induction Condition 1 (❶) and Condition 2 (❷) for Pax7 (magenta) and Pitx2 (green). Higher magnification views as insets in (A’, B’, C’). White arrow heads indicate Pax7+/Pitx2+ cells; open arrow heads indicate Pax7+/Pitx2- cells, yellow arrowheads indicate Pitx2+ myonuclei. **(D)** Percentage of Pitx2+/Pax7+ cells over total Pax7+ cells in EOM from induction Condition 1 (❶) and Condition 2 (❷) (WT, KO n=3 each). Scale bars: 100μm (A-C). Error bars represent SEM. ***P<0.001, ****P<0.0001.

**Supplementary Figure 6.**
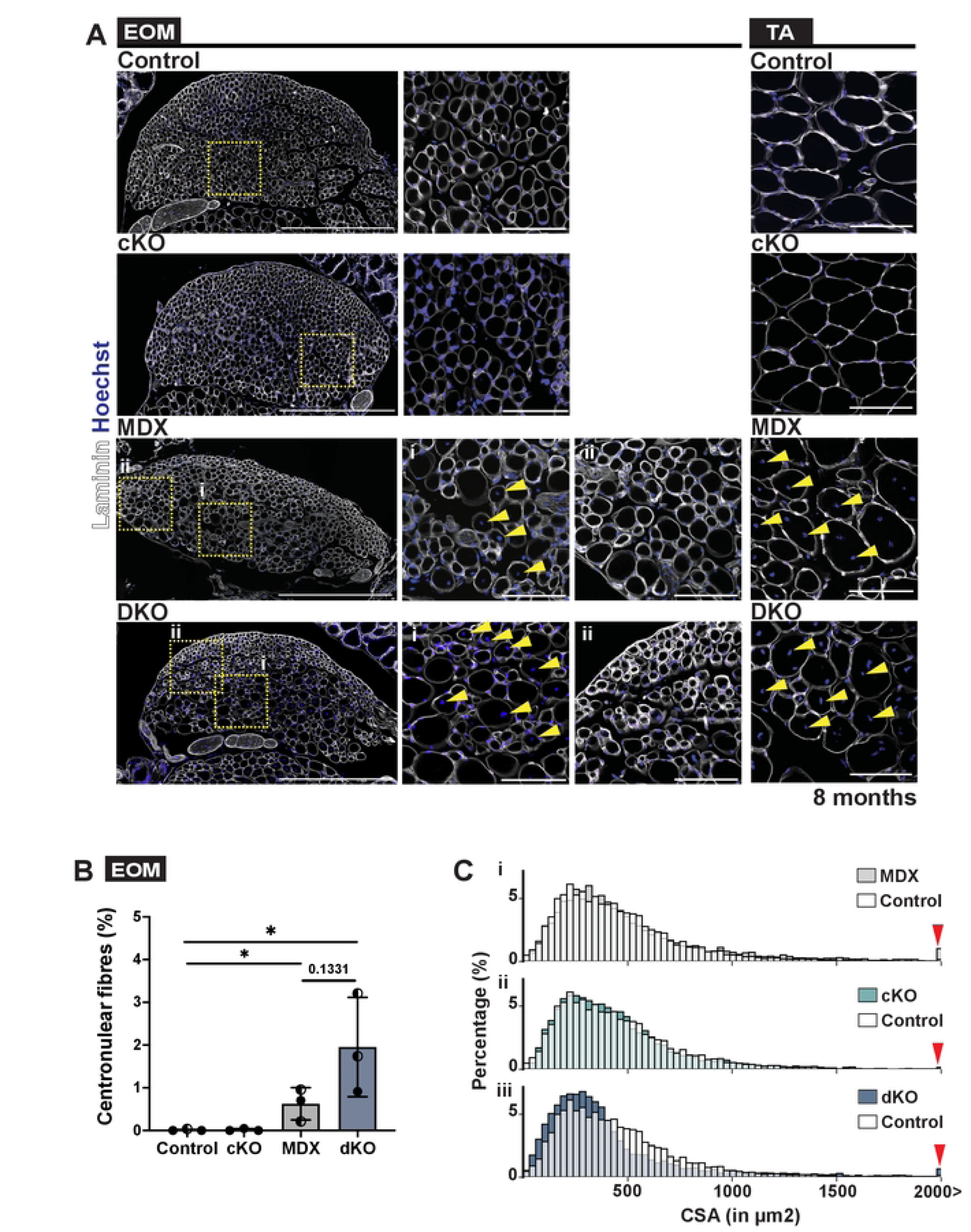
Early postnatal deletion of *Pitx2* in EOM MuSCs affects myofibre size in *mdx* mice. **(A)** Immunostaining of EOM sections and TA from Control (*Pitx2^flox/flox^* or *Pitx2^flox/+^*), MDX and MDX Pitx2cKO (dKO) mice at 8 months of age for laminin (white). Right panels (i, ii), higher magnification views of the area delimited with dots. (i) EOM regions containing central myonuclei, (ii) nomal EOM regions. Yellow arrow heads indicate central myonuclei. **(B)** Number of centrally nucleated myofibres in EOM from immunostaining in (A). **(C)** Distribution of EOM myofibre size from immunostaining in (A) (Control, cKO and MDX, n=3; dKO, n=4). Red arrowheads indicate fibres over 2000 μm observed in MDX and dKO. Scale bars: (A) 1000 μ m in lower magnification, 100 μ m in higher magnification views. Error bars represent SEM. *P<0.05.

